# Causal analysis of rock–biodiversity relationships over the global Phanerozoic challenges rock record bias and common cause hypothesis

**DOI:** 10.64898/2026.10.22.761025

**Authors:** Alexis Rojas, Gregor Mathes, Andreas Futschik

## Abstract

Sampled marine biodiversity covaries strongly with sedimentary rock quantity over the Phanerozoic, but the respective contributions of different potential causes remain unsettled. Although regional studies show that rock-record bias accounts for more variation in sampled marine biodiversity than common-cause mechanisms, evaluating this relationship across the global Phanerozoic record remains difficult because fossil databases are inherently biased toward fossiliferous strata. To address this limitation, we compiled literature-derived measures of global marine biodiversity and sedimentary strata, along with an existing sea-level curve. Using an information-theoretic causal framework, we then evaluated the relative contributions of rock-record bias and common-cause hypotheses to Phanerozoic-scale variation in sampled marine biodiversity. Our results show a strong contemporaneous causality of marine strata on sampled biodiversity, indicating that rock record bias detected at regional scales extends to the global scale. However, neither rock record bias nor common-cause mechanisms accounts for most Phanerozoic-scale patterns in sampled biodiversity, and additional unmeasured factors likely contribute. Finally, we find tentative evidence that synergistic causality between geologically mediated sampling biases and forces shaping the underlying biological signal contributes to global Phanerozoic patterns in sampled biodiversity.

## 1. INTRODUCTION

Sampled marine diversity strongly covaries with quantity of preserved sedimentary rock (Dunhill, 2011; Hannisdal and Peters, 2011). Two main hypotheses have been proposed to explain this pattern. The rock record bias postulates that temporal variation in rock availability influences observed biodiversity patterns through differences in preservation and sampling (Raup, 1972). In contrast, the common-cause hypothesis proposes that geological processes affecting the sedimentary record also act as macroevolutionary forces shaping the underlying biological signal (Peters, 2005). Regional studies have shown that rock-record bias accounts for more variation in sampled marine biodiversity than common-cause mechanisms (Dunhill et al., 2014; Na et al., 2023). However, their respective contributions to global Phanerozoic variations in sampled marine biodiversity remain unresolved. This uncertainty stems from two major challenges. First, fossil databases are inherently biased toward fossiliferous strata, making it difficult to distinguish genuine fossil absence from gaps in the sedimentary record at the global scale (Na et al., 2023). Second, correlation-based approaches commonly used in paleontology cannot distinguish causal relationships from statistical associations induced by common causes (Runge et al., 2023).

To address these issues, we compiled literature-derived measures of global marine sedimentary strata and sampled biodiversity, together with an existing sea-level curve. Using the SURD framework, we decomposed the causal information that observed variables provide about future sampled biodiversity into unique, redundant, and synergistic components, while quantifying causality leakage due to unobserved variables. We also formalized the two competing hypotheses within an explicit causal framework to clarify their underlying assumptions and provide a basis for their systematic evaluation.

### Rock record bias

The rock record bias (sampling bias *sensu* Raup, 1972) postulates a strong, potentially causal influence of preserved sedimentary rock on sampled Phanerozoic marine diversity, acting primarily through its influence on sample size (Alroy et al., 2008). It is consistent with the notion that the benthic marine fossil record is linked to the preservation of habitable facies in the sedimentary record (Holland et al., 2025). The rock record bias represents a direct-effect mechanism in which rock quantity is treated as primary driver of sampled biodiversity. Despite the methods developed to correct for collection biases (e.g., Alroy et al., 2008; Alroy, 2010; Close et al., 2018), estimating rock quantity from fossiliferous strata continues to obscure interpretations of rock-biodiversity relationships (Peters, 2005; Na et al., 2023). Because sea-level fluctuations influence the relative proportions of shallow- and deep-water continental-shelf deposits preserved at large spatial scales (Smith, 2001), rock quantity could mediate the influence of sea-level change on sampled biodiversity (Figure 1A).

**Figure 1.**
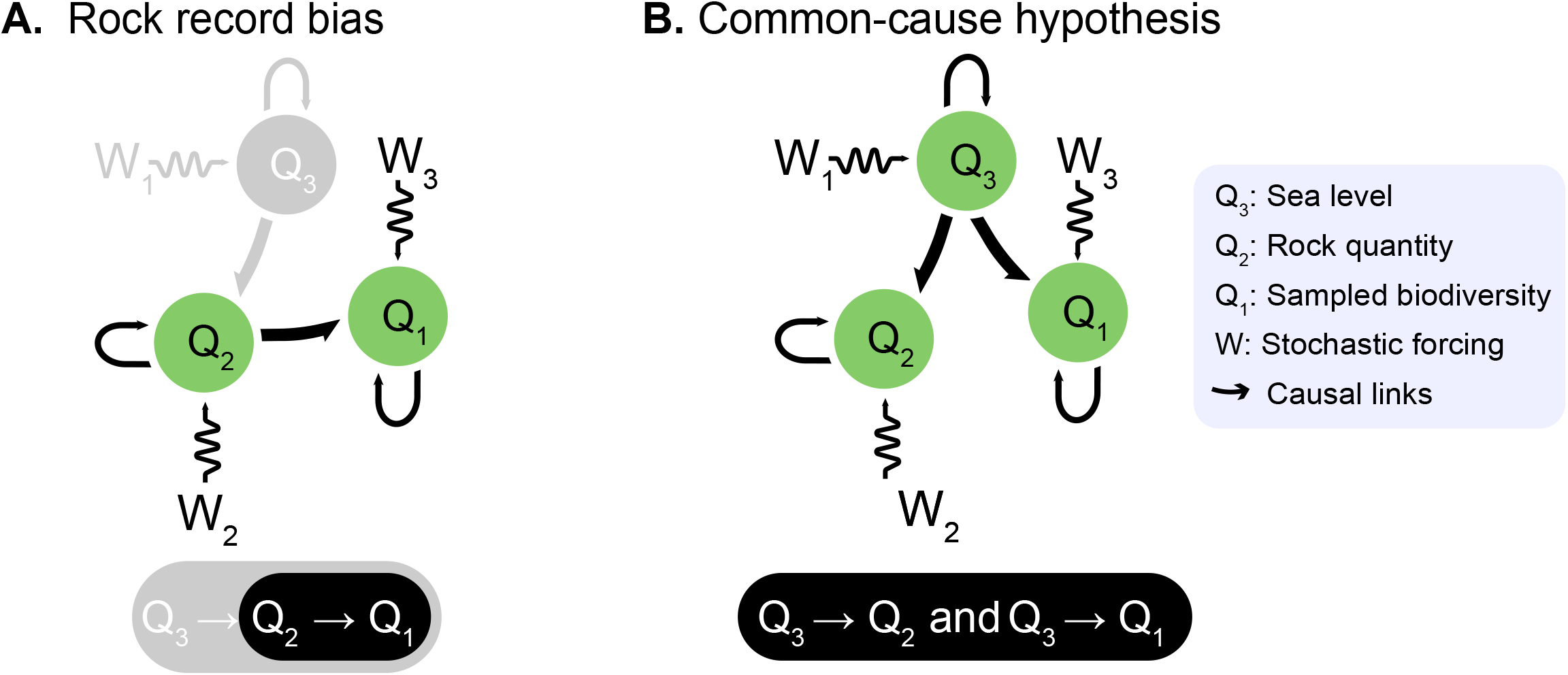
Schematic representation of the functional dependencies among variables in the competing hypotheses. **A.**Rock record bias.**B**. Common-cause hypothesis.

### Common-cause hypothesis

The common-cause hypothesis postulates that variations in rock quantity represent fluctuations in sea-level that also directly modulate true marine (*sensu* Peters, 2005), acting primarily by changing the area of benthic marine habitat (Bambach, 1999; Foote, 2000). It is consistent with the notion that processes generating variability in the sedimentary rock record, such as sea-level change and plate tectonics, may have been also important macroevolutionary forces (Zaffos et al., 2017; Mathes et al., 2021). The common-cause hypothesis has been invoked to explain true biodiversity and therefore focuses on the role of evolutionary processes.

Common-cause mechanisms have also been proposed to explain similarities between patterns of evolutionary rates and the sedimentary record (e.g., Bambach, 1999; Foote, 2000; Peters, 2005). This hypothesis assumes that sampled biodiversity primarily reflects variation in the underlying biological signal, thereby accounting for the observed congruence between rock quantity and sampled biodiversity. If sea level causally affects rock quantity and biodiversity, it may act as a confounder (Cochran, 1957), inducing a spurious association between the fossil and rock records (Figure 1B).

## 2. DATA AND METHODS

### Bibliographic analysis

We compiled literature-derived proxies for global marine strata, fossiliferous strata, and marine biodiversity by quantifying literature associated with sedimentary rocks across a defined set of depositional environments and geological bins (Figure 2 A-D). Documents were compiled from the Web of Science (WoS) Core Collection using a search string designed to retrieve those with content associated exclusively to individual temporal bins (Data S1). This strategy was adapted to search for taxa representing well-preserved benthic marine invertebrates (Kocsis et al., 2018) and for rocks deposited in the corresponding marine environments. Rock names were obtained from the Paleobiology database (PaleoDB) *lithology* field of collections in which target taxa are reported (Peters and McClennen, 2016). Depositional environments follow the PaleoDB *Depositional environment tip sheet* (Uhen et al., 2026). Documents were collected separately for each interval in the Ten Million Year Bins timescale. Publication counts from documents sets retrieved via rock-related search are used as proxies proxy for sedimentary strata. Target genera were sourced from the PaleoDB, and their occurrences within the document sets were identified using exact string-matching (McLevey and McIlroy-Young, 2017) (Notebook S1).

**Figure 2.**
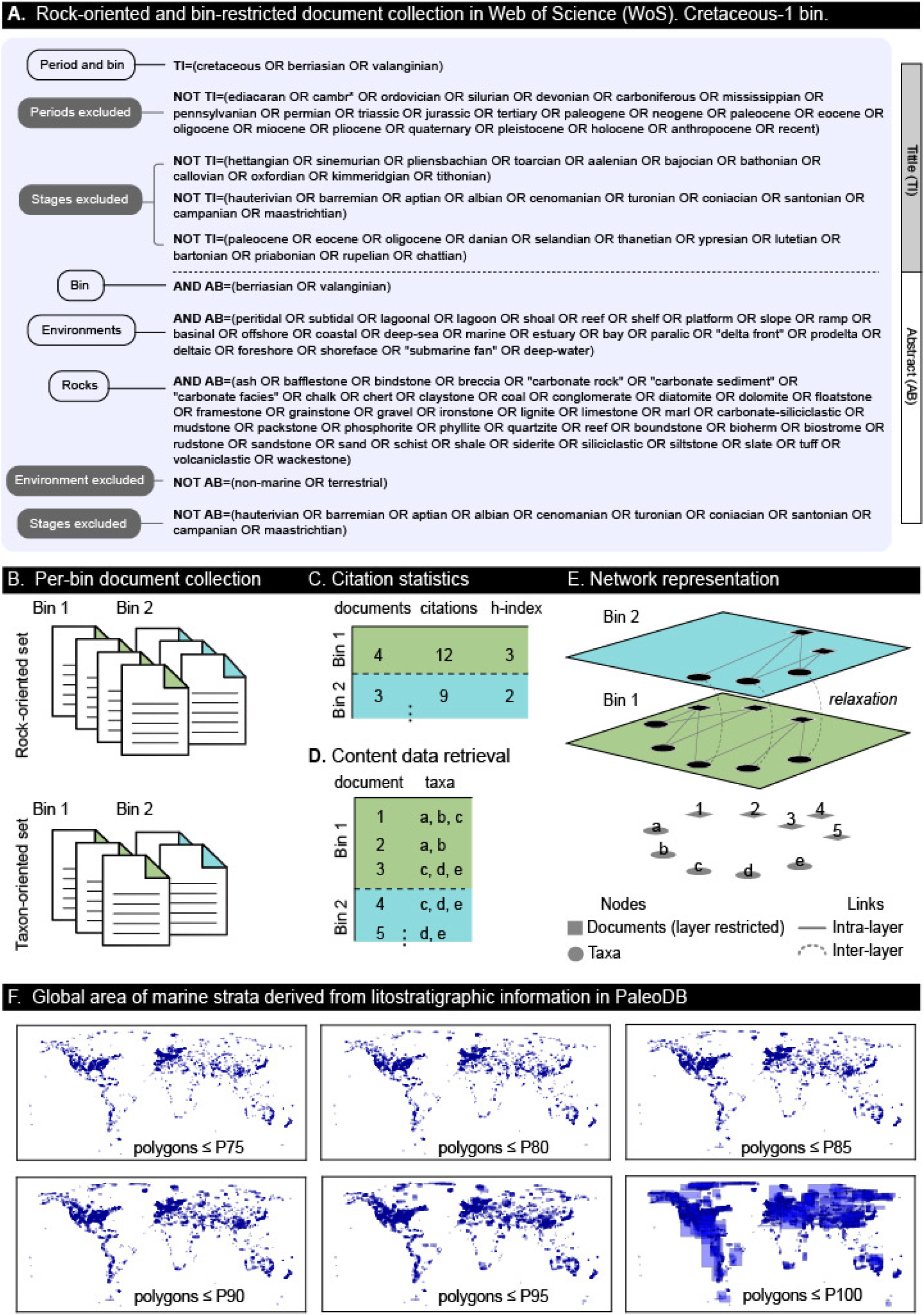
Workflow.**A.**Search string for bin-restricted document collection. **B**. Document collection. Environment-, taxon-, and rock-oriented searches were performed. **C**. Marine strata proxy. It is the number of documents recovered from rock-oriented searches. **D**. Sampled biodiversity proxy. It is the number of genera recovered from rock-oriented searches. **E**. Network representation. **F**. Global area of marine strata. Polygons representing formations filtered using percentile-based thresholds for both dimensions.

The assembled dataset [bin × document × taxon] has a structure similar to the network described by Rojas et al. (2021). We therefore use a network framework to assess whether it captures the known temporal distribution and relative representation of the different taxonomic groups. Specifically, we constructed a multilayer network (Figure 2E; Data S2), where layers represent ordered temporal bins (Rojas et al., 2022b). Consistent with the bin-restricted collection, documents are represented as layer-dependent nodes. Within each layer, documents and taxa are connected by intra-layer unweighted links (Rojas et al., 2017, 2022a). Inter-layer connections are generated via relaxation, linking identical taxa across successive bins (Eriksson et al., 2021, 2022). The map equation (Edler et al., 2017) is used to measure normalized flow, describing how a random walker moves through individual nodes in the network.

### GIS analysis

We used marine strata from the PaleoDB to generate an independent, broad comparative estimate of rock record quantity and assess whether our proxy captures similar temporal patterns. Polygons representing lithostratigraphic units were generated using an equal-area Mollweide projection (Snyder, 1987), quantified by their longitudinal and latitudinal extents, and filtered using percentile-based thresholds for both dimensions. Global maps and area statistics were used to assess the geographic distribution of the filtered units and the effects of progressively excluding the largest polygons. Filtered polygons were assigned to temporal bins based on median age, and polygons whose durations exceeded the corresponding bin duration by specified percentages were excluded (Figure 2F).

### Causal analysis

We employed the Synergistic-Unique-Redundant Decomposition (SURD) framework (Martínez-Sánchez et al., 2024) to assess forward-in-time and contemporaneous causal relationships between marine strata and sampled biodiversity. SURD decomposes the causal information that source variables provide about a target variable into unique (U), redundant (R), and synergistic (S) components. It quantifies the causal influence of the vector of observations *Q* to the future of the target variable *Q*^+^ = *Q*_j_ (*t* + ΔT), where *Q*_j_ is one of the components of *Q* and ΔT > 0 is an arbitrary time lag. For a subset *Q*_*i*_ = [*Q*_*i*1_, *Q*_*i*2_, …] of *Q*, redundant causality from *Q*_*i*_ to *Q*_*j*_^+^ is information shared among the components of *Q*_*i*_, unique causality is information from *Q*_*i*_ that cannot be obtained from any other variable, and synergistic causality arises from the joint effect of variables in *Q*_*i*_. Causal components are normalized using the mutual information between all observed variables and *Q*_*j*_^+^. Causality from unobserved variables that influence *Q*_*j*_^+^ but are not included in *Q* is quantified as information leak (ΔI_leak_).

## 3. RESULTS

### Literature-derived proxies for marine biodiversity and strata

The assembled dataset comprises 9013 documents containing 3971 occurrences representing 2493 genera (Data S1; Figure 3A). The per-bin number of genera recovered through rock-oriented searches serves as a proxy for sampled biodiversity and shows a significant positive correlation with genus occurrences in PaleoDB (Spearman’s ρ = 0.50, p < 0.001). This correlation is similar for the total genera obtained from combined rock-, taxon-, and environment-oriented searches (Spearman’s ρ = 0.54, p < 0.001; Figure 3B). Similarly, the per-bin number of documents recovered through rock-oriented searches serves as a proxy for global marine strata, independent of fossil content. It shows a significant positive correlation with the global area of marine strata reconstructed from lithostratigraphic units in PaleoDB (Spearman’s ρ = 0.66, p < 0.001; Figure 3C). The subset of documents reporting genera from the target invertebrates serves as a proxy for fossiliferous strata and shows a significant positive correlation with the number of formations in PaleoDB (Spearman’s ρ = 0.53, p < 0.001; Figure 3D).

**Figure 3.**
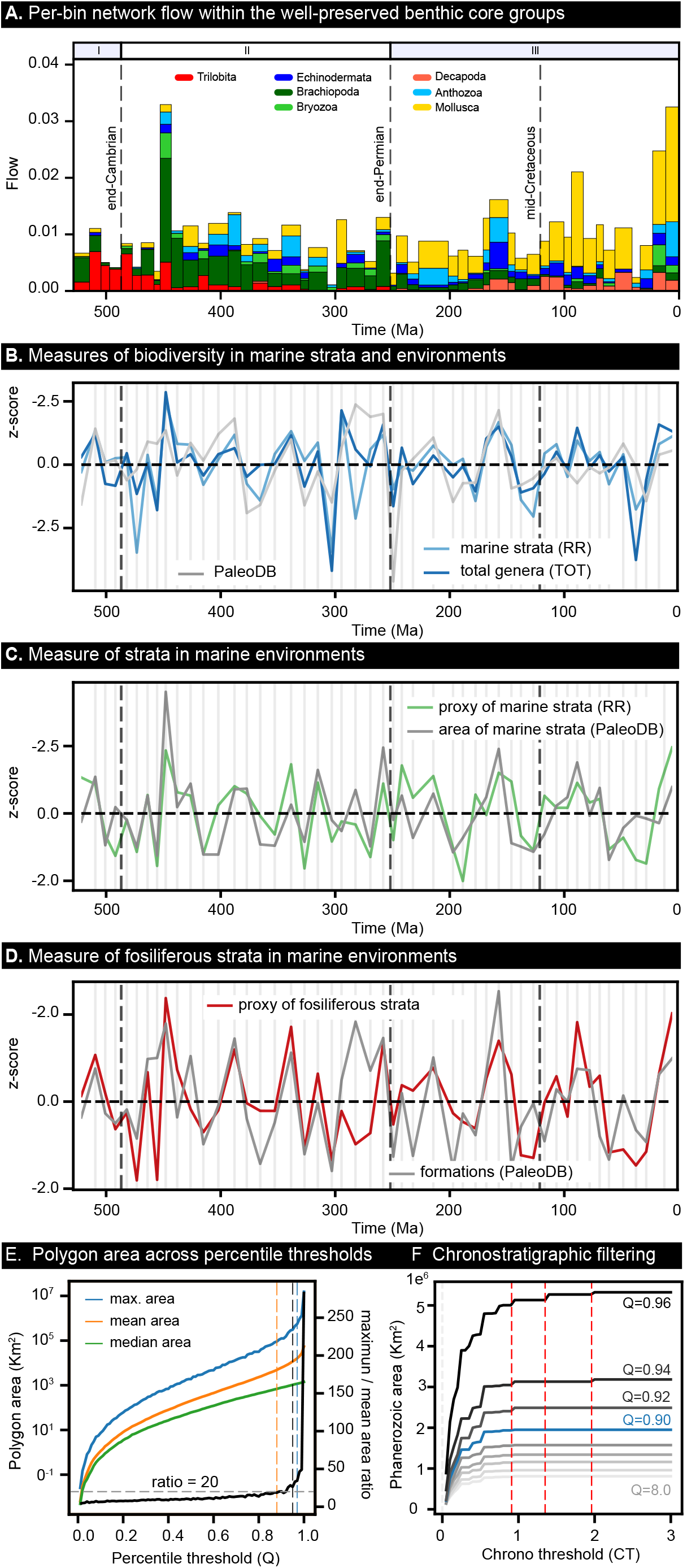
**A**. Phanerozoic distribution of marine invertebrates in the assembled dataset. I-IV indicate intervals dominated by each global mega-assemblage. **B**. Sampled biodiversity proxy. Genera from combined searches (TOT) and PaleoDB are shown for reference. **C**. Marine strata proxy. Global area of marine strata is shown for reference. **D**. Fossiliferous strata proxy. Subset of documents reporting genera from the target invertebrates. PaleoDB formations shown for reference. All variables are log-transformed. Long-term trends are estimated using locally weighted regression (smoothing parameter = 0.3). Detrended time series are residuals from LOESS fits to log-transformed observations, standardized as z-scores. **E**. Global area of marine strata across percentile thresholds. **F**. Effect of chronostratigraphic filtering on the Phanerozoic area of marine strata.

### Global marine area

Global maps of polygons representing geological formations, (Figure S2), polygon-area statistics, and knee-point analysis together indicate that percentile thresholds of approximately 0.80–0.90 yield stable spatial patterns while limiting the influence of exceptionally large polygons (Figure 3 E). In addition, the total Phanerozoic area retained is robust to chronostratigraphic filtering (CT), with the same polygons retained when durations up to 50% longer than the bin duration are allowed (Figure 3 F). Therefore, the per-bin area summed across all filtered polygons (Q = 0.9, CT = 0.5) is used as a broad estimate of rock quantity. It shows a significant positive correlation with our proxy for marine strata (Spearman’s ρ = 0.66, p < 0.001).

### Causal relationship between marine strata and sampled biodiversity

Sampled biodiversity is primarily influenced by the unique contemporaneous causality from marine strata (U_5→1_; Figure 4). followed by synergistic causalities involving marine strata, sea level and sampled biodiversity that propagate forward-in-time (S_23→1_, S_123→1_). The only unique causality on marine strata is the forward-in-time influence of sea level (U_3→2_), whereas the only unique causality on sea level is self-induced (U_3→3_). Synergistic causalities propagating forward in time and affecting marine strata and sea level at magnitudes comparable to those of the observed unique causalities could reflect complex causal pathways, artifacts of the assembled dataset, or noise arising from the limited length of the time series (Martínez-Sánchez et al., 2024). In addition, the high contemporaneous causality from sampled biodiversity to marine strata (U_4→2_) likely reflects an orientation artifact, as its direction is consistent with the high-magnitude unique causality from marine strata to sampled biodiversity (U_5→1_) and with the established mechanism underlying rock record bias. Remarkably, the causality leak (ΔI_leak_) exceeds 55% in forward-in-time and contemporaneous analyses.

**Figure 4.**
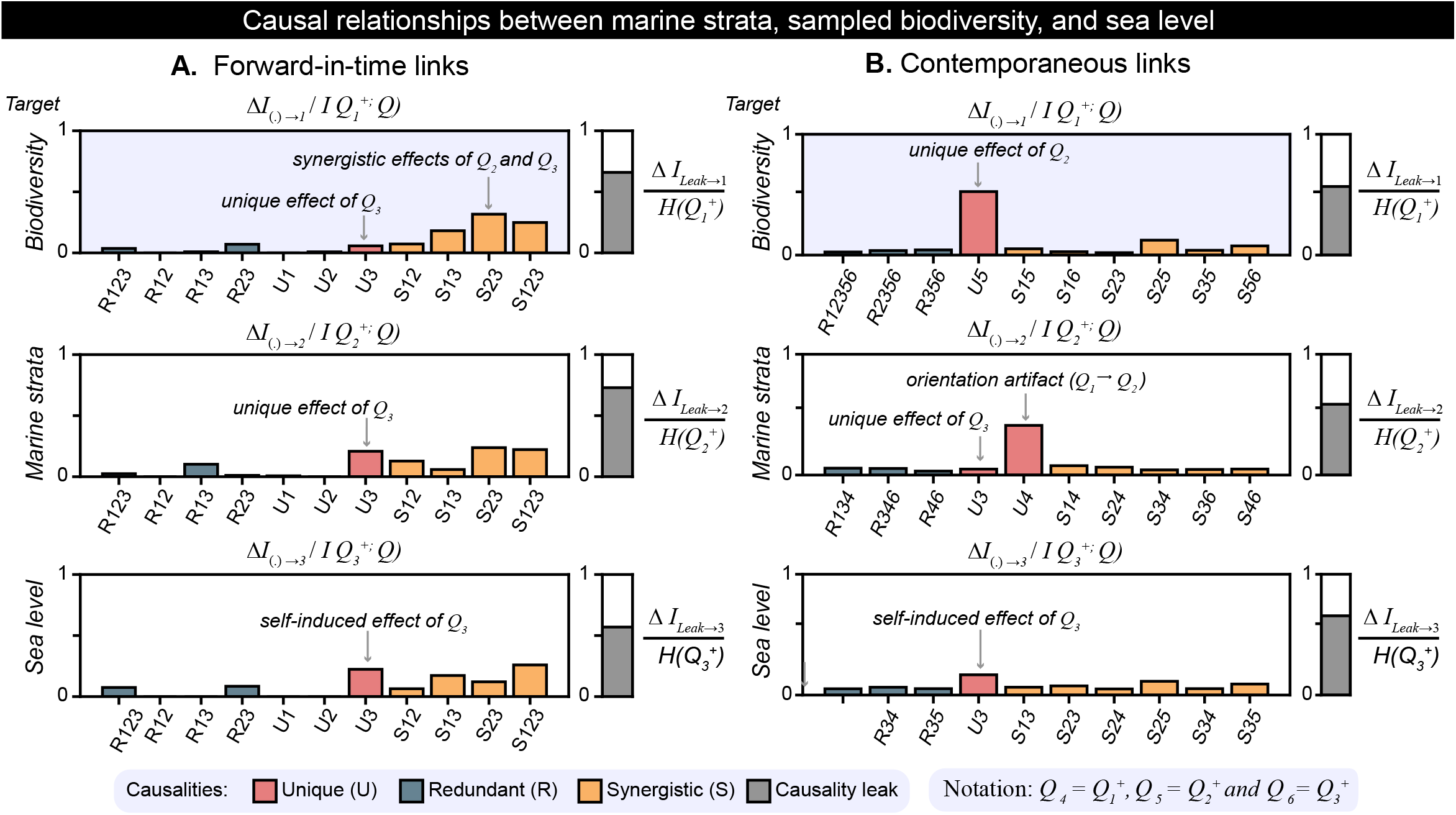
SURD results.**A.**Forward-in-time causal analysis. **B**. Contemporaneous causal analysis. **C–D**. Causal components: unique (U), redundant (R), and synergistic (S). *Q*^+^, target variable; *Q* set of observations; ΔT =1 for the forward-in-time analysis and ΔT = 0 for the contemporaneous analysis; ΔI_leak_, causality leak.

## 4. DISCUSSION

The assembled dataset captures the chronostratigraphic distribution of well-preserved benthic marine invertebrates and is consistent with major extinction events and the mega-assemblages that successively dominated the Phanerozoic oceans (Kröger and Rojas, 2026). The agreement between our literature-derived measures and independent PaleoDB records supports their use as proxies for marine strata and sampled biodiversity across the Phanerozoic. In particular, the correspondence between our proxy for marine strata and independent measures based on lithostratigraphic information indicates that the proxy broadly captures variation in rock quantity through time while remaining independent of fossil content. Because WoS searches a scholarly metadata and citation graph and automatically expands queries, its standard search is unsuitable for time-restricted content retrieval. Our approach overcomes this limitation by restricting searches to titles and abstracts and specifying taxa, temporal intervals, rocks and sedimentary environments to be included and excluded.

SURD analysis indicates a strong contemporaneous dependence of sampled biodiversity on marine strata, consistent with the hypothesized causal influence of preserved sedimentary rock on sampled marine diversity underlying the rock record bias (Alroy et al., 2008). It also provides tentative evidence for forward-in-time synergistic causality from sea level and marine strata to sampled biodiversity, consistent with a joint influence of geologically mediated sampling biases and forces shaping the underlying biological signal on global Phanerozoic patterns in sampled biodiversity. Given the relatively short time series (50 bins), evidence for other weak causalities should be interpreted cautiously, as these signals may reflect artifacts (Martínez-Sánchez et al., 2024). We could not compile the dataset at stage-level resolution because searches yielded no documents for some stages. Nevertheless, SURD results indicate that neither rock-record bias nor a common-cause mechanism involving sea level as a physical forcing factor accounts for most of the global Phanerozoic-scale variation in marine sampled biodiversity.

## 5. CONCLUSIONS

Our literature-derived proxies capture the temporal distribution of benthic marine invertebrates and the marine sedimentary record throughout the Phanerozoic. SURD analysis provides evidence for a strong contemporaneous causality from marine strata to sampled biodiversity, indicating that the rock-record bias previously documented at regional scales extends to the global Phanerozoic. In addition, the results provide tentative evidence for forward-in-time synergistic causality from sea level and marine strata to sampled biodiversity, suggesting that interactions between geologically mediated sampling biases and forces shaping the underlying biological signal may contribute to observed Phanerozoic patterns in biodiversity. However, neither the rock-record bias nor a common-cause mechanism involving sea level accounts for most Phanerozoic-scale variation in sampled marine biodiversity. Instead, additional unmeasured factors likely contribute to the observed congruence between the marine rock and fossil records throughout the global Phanerozoic.

## ACKNOWLEDGMENTS

This work was supported by the Johannes Kepler University Linz, Linz Institute of Transformative Change (LIFT_C).

## Supplementary Material

**Data S1**. Search strings and citation statistics for the document collections.

**Data S2**. Assembled dataset in multilayer format.

**Figure S1.**
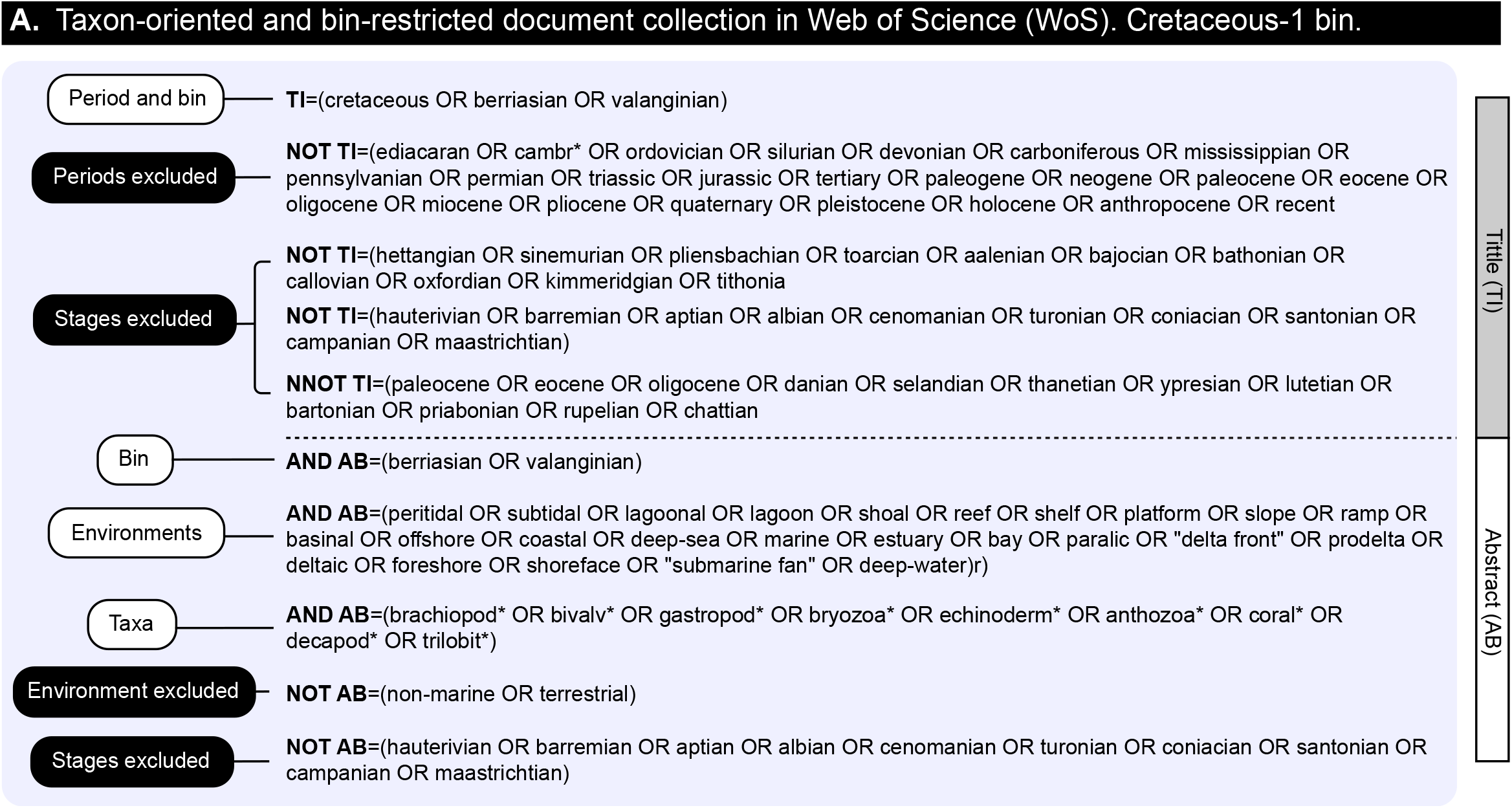
Search string for taxon-oriented document collection.

**Figure S2.**
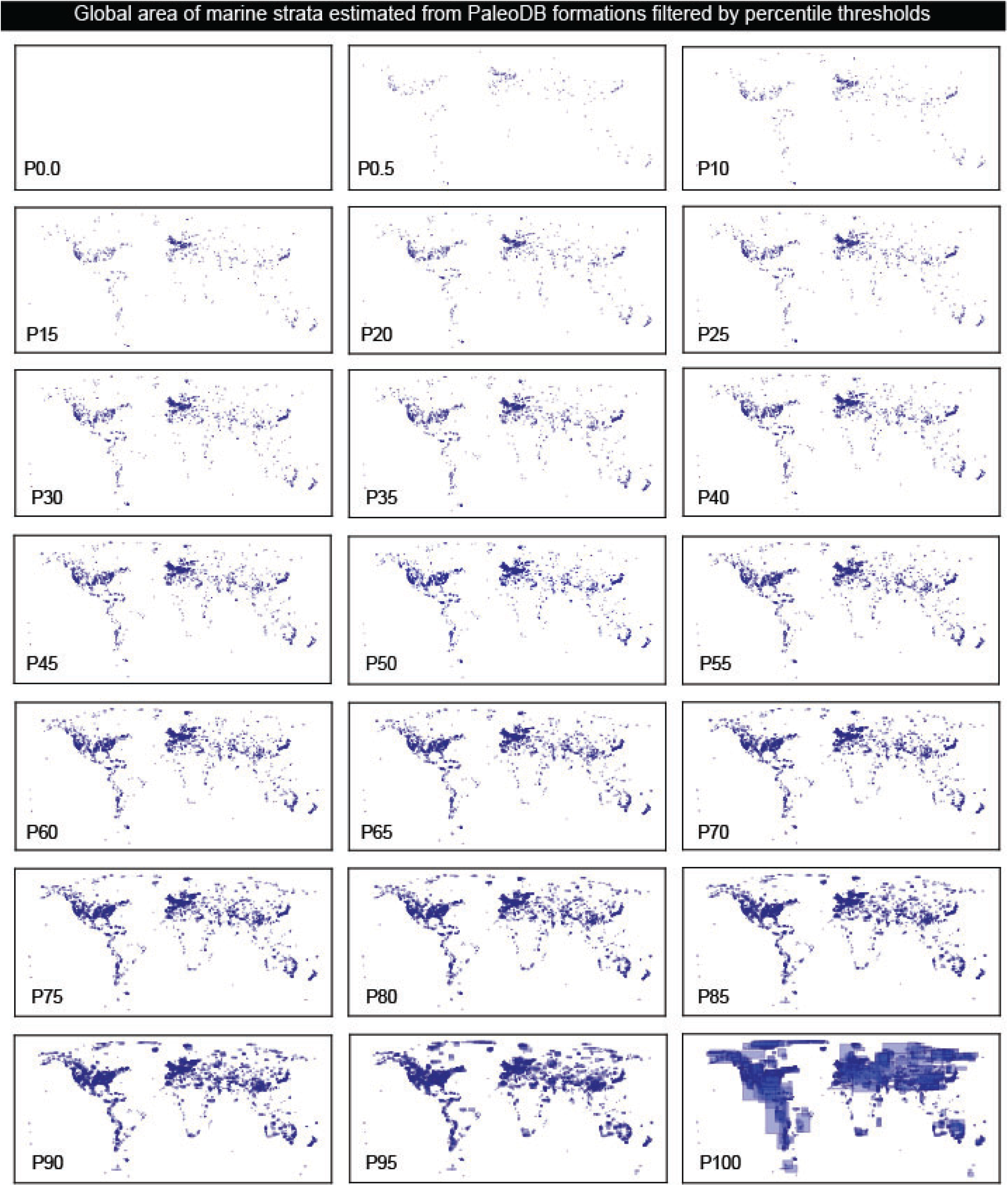
Global area of marine strata across percentile

